# Network analysis of Differentially Expressed Genes (DEGs) identified in zebrafish after infection with Spring viremia of carp virus (SVCV) – an *in silico* approach

**DOI:** 10.1101/2022.03.01.482441

**Authors:** Praveen Kumar Guttula, Mohd Asharf Rather

**Affiliations:** Department of Clinical Hematology, Tata Medical Center, Kolkata 700160, India; Division of Fish Genetics and Biotechnology, Faculty of Fisheries Rangil, Ganderbal, SKUAST-Kashmir, India

**Keywords:** Spring viremia of carp virus (SVCV), RNA sequencing, Kyoto Encyclopedia of Genes and Genomes, transcriptome analysis

## Abstract

Spring viremia of carp virus (SVCV) is a virus that belongs to family of spring viremia of carp (SVC) and frequently causes hemorrhagic symptoms in several types of cyprinids and causes severe economic and environmental losses. Therefore, the mechanism of the infection is not clearly understood. In this study, zebrafish was employed as the infection model to explore the pathogenesis of SVCV. 4 groups of zebrafish tissues were set and RNA sequencing (RNA-Seq) technology was employed to analyze the differentially expressed genes (DEGs) after SVCV-infection. A total of 360,971,498 clean reads were obtained from samples, 382 DEGs in the brain and 926 DEGs in the spleen were identified. These DEGs were annotated into three ontologies after gene ontology (GO) enrichment analysis. The Kyoto Encyclopedia of Genes and Genomes (KEGG) pathway analysis showed that these DEGs were primarily related to Influenza. A pathway and Herpes simplex infection pathway in brain and Tuberculosis and Toxoplasmosis pathways in spleen, and all of these pathways may be involved in response to pathogen invasion. The transcriptome analysis results demonstrated changes and tissue-specific influences caused by SVCV in vivo, which provided us with more information to understand the complex relationships between SVCV and its host.

## 1. Introduction

Over the years, fish farming has been adversely affected by viral infections such as rhabdoviruses that are considered one of the most devastating diseases in the fishery. For instance, a seasonal disease caused in Central Europe by the spring viremia carp virus (SVCV) negatively affects the warm-water cyprinid fish farms [1]. Although, there are multiple interventions to prevent viral infections in fish, among the DNA vaccines theoretically seems might be used to combat the fish pathogens. However, practically these DNA vaccines often fail to fulfill expectations in preventing these viral infections in fish. For example, DNA vaccines are effective for novirhabdoviruses that produce a non viral NV protein but are found not effective against spring viremia carp virus (SVCV), which belongs to as novirhabdoviruses family [2]. Hence, studying the difference in fish infection/vaccination between these two rhabdoviral models has become an exciting field of research.

Moreover, there is a need for a better understanding of basic knowledge on fish immune responses that helps in improving or developing new DNA vaccine against novirhabdoviruses that are injectable to more practical oral immunizations of fish [2, 3]. The use of microarrays could greatly expand this elemental knowledge as well as aid in the development of other fish DNA vaccines [4]. Due to the lack of known genes that are associated with survival during viral infections, research on the possibility of using drugs to prevent fish viral infections has not been conducted. Because of its cyprinid status and genome sequence and genetics, the zebrafish (*Danio rerio*) is an ideal model to study SVCV infections and zebrafish have no infectious diseases in their natural state [5]. Furthermore, as the zebrafish are vulnerable to both the rhabdoviruses that lack the NV gene and infect warm-water fish, such as SVCV [6], and novirhabdoviruses that encode for NV and influence cold-water fish, such as infectious hematopoietic necrosis (IHNV), snakehead rhabdoviruses (SHRV) [7], and viral hemorrhagic septicemia (VHSV) [8], zebrafishes can be utilized for comparative analysis of the NV-coding and the NV-lacking fish rhabdoviral group. VHSV was used in a prior zebrafish microarray investigation because it has been shown to cause a natural mode of infection through immersion in both adults as well as larvae, and an effective immunization had been recorded [9]. However, successful VHSV infection requires zebrafish acclimatization to temperatures lower than their ideal 14 °C, thereby making the effects of SVCV infections at 24°C easier to understand without the distraction of probable low temperatures.

Microarrays have not yet been used to analyze fish transcriptome alterations following the immunization with fish rhabdoviruses lacking the NV gene, such as SVCV. Wide-genome microarrays, on the other hand, have been used to estimate the expression profiling of numerous fish genes through infections with Novirhabdoviruses such as Hiramerhabdovirus (HRV), VHSV, and IHNV [10]. Sequencing technologies, such as 454-pyrosequencing, have recently been used in the VHSV/turbot model [11]. The maximum number of significantly expressed fish genes was observed 2-3 days after infection in the majority of the investigations listed above. In light of these findings, the current research is concentrated on the response of zebrafish to acute infection (2 days) and then compared with those of survivor fish of about 30-days old.

Recent advancements in bioinformatics methodologies [12, 13] have provided novel and robust tools for the biomarker identification of various diseases [14, 15]. Network modeling techniques are capable of integrating and improving the possibility of disease state complexity and its causative factors. Network analysis approaches also provide unique and practical strategies for the prediction of the diseases at an early stage and drug designing [16]. As a result, we decided to look into the host genes that were most associated with the SVCV/zebrafish infection model to see if there were any potential medications to prevent the disease hypothetically. Thus, in the current study, an integrative bioinformatics approach was applied to gene expression data, from infected and uninfected zebrafish, to identify hub genes, and decipher the molecular mechanism of viral infection by network analysis. The differentially expressed genes (DEGs) were analyzed using the gene ontology, pathway analysis and network analysis, and corresponding hub genes were identified. These findings could help researchers to create new drug-based preventative techniques and expand the use of the zebrafish/SVCV model to explore additional vertebrate viral infections.

## 2. Materials and Methods

### 2.1. Screening of Differentially Expressed Genes (DEGs)

The dataset with GEO Id GSE63133 [17] from Gene expression omnibus (GEO) database [18] was selected for the analysis. The head kidney and spleen tissues of the SVCV infected zebrafish and uninfected zebrafish were used in the above dataset. The DEGs which were present as a supplementary data in the above study were retrieved and selected for prediction of the biomarkers. The log2FC values >=2 and <=-2 were used to select the DEGs between the infected and uninfected groups.

### 2.2. Functional annotation and pathway enrichment analysis

Retrieved DEGs were submitted to DAVID (Database for Annotation, Visualization, and Integrated Discovery) (https://david.ncifcrf.gov/) functional annotation tool [19]. The Gene Ontology (GO) and KEGG pathway analysis was performed to identify the genes that are being enriched with biological process (BP), molecular function (MF), cellular component (CC) and biological pathways. The enrichment analysis was performed using default settings and with p < 0.05.

### 2.3. Protein-Protein Interaction Network and hub gene prediction

A set of DEGs were submitted to STRING (Search Tool for the Retrieval of Interacting Genes/Proteins) database (https://string-db.org/) [20], which analyzes the various relationships among the targets through a graphical interface by creating a protein-protein interaction network. The STRING interactions.tsv file was imported into Cytoscape software (version 3.8.2) [21]. Various Cytoscape plugins like MCODE [22], Centiscape 2.2 and CytoHubba were used to analyze the complex biological PPI network. MCODE [22] was used to divide the whole interaction network into all possible clusters (highly interconnected regions) with the default parameters like loops, haircut, fluff, node score cutoff-0.2, k-core-2 and max depth of 100 were selected. Further Centiscape analysis was applied to each MCODE derived cluster to study the various centralities measures (degree, closeness and betweenness) among the interconnected nodes within the clusters. Accordingly the high scores of degree, betweenness and closeness, the important regulatory nodes were screened out separately for each cluster. Cytoscape plugin CytoHubba was used to analyze the complex network in order to evaluate the top ten hub nodes through maximal clique centrality (MCC) scoring method in the PPI network.

## 3. Results and Discussion

### 3.1. Screening of DEGs

Variation between the gene expressions was studied between the infected and uninfected zebrafish. The screening of the DEGs was done based on the method described in the materials and methods section. The results showed that a total 227 genes were expressed differentially between the infected and uninfected zebrafish. Out of 227 genes 90 genes were up regulated and 137 genes were down regulated. The lists of genes were provided in the supplementary data as Table S1 (**Page No 1-4**).

### 3.2. Functional annotation and pathway analysis

The functional annotation study revealed the involvement of six biological processes, two cellular components, seven molecular functions and six KEGG pathways with our significant genes. The associated GO terms in biological process were cellular response to estrogen stimulus (GO:0071391) with p value= 5.54E-05, lipid transport (GO:0006869) with p value=0.001069, complement activation (GO:0006956) with p value=0.014337, blood coagulation, fibrin clot formation (GO:0072378) with p value=0.014337, aromatic amino acid family metabolic process (GO:0009072) with p value=0.034072, proteolysis (GO:0006508) with p value=0.065149 etc, in cellular components were extracellular region (GO:0005576) with p value= 4.66E-09 and extracellular space (GO:0005615) with p value= 1.46E-08; in molecular function were endopeptidase inhibitor activity (GO:0004866) with p value=7.02E-08, serine-type endopeptidase inhibitor activity (GO:0004867) with p value=4.91E-06, scavenger receptor activity (GO:0005044) with p value=5.91E-04, lipid transporter activity (GO:0005319) with p value= 0.001086, serine-type endopeptidase activity (GO:0004252) with p value= 0.0037, polysaccharide binding (GO:0030247) with p value=0.051412, cysteine-type endopeptidase inhibitor activity (GO:0004869) with p value=0.054537 etc. The KEGG pathways such as metabolic pathways (dre01100) with p value = 0.004386, cysteine and methionine metabolism (dre00270) with p value= 0.007229, biosynthesis of antibiotics (dre01130) with p value = 0.029217, ubiquinone and other terpenoid-quinone biosynthesis (dre00130) with p value = 0.029235, phenylalanine metabolism (dre00360) with p value = 0.046389, tyrosine metabolism (dre00350) with p value = 0.093463 etc were involved with these target genes. These results were shown graphically in **Figure 1**.

**Figure 1:**
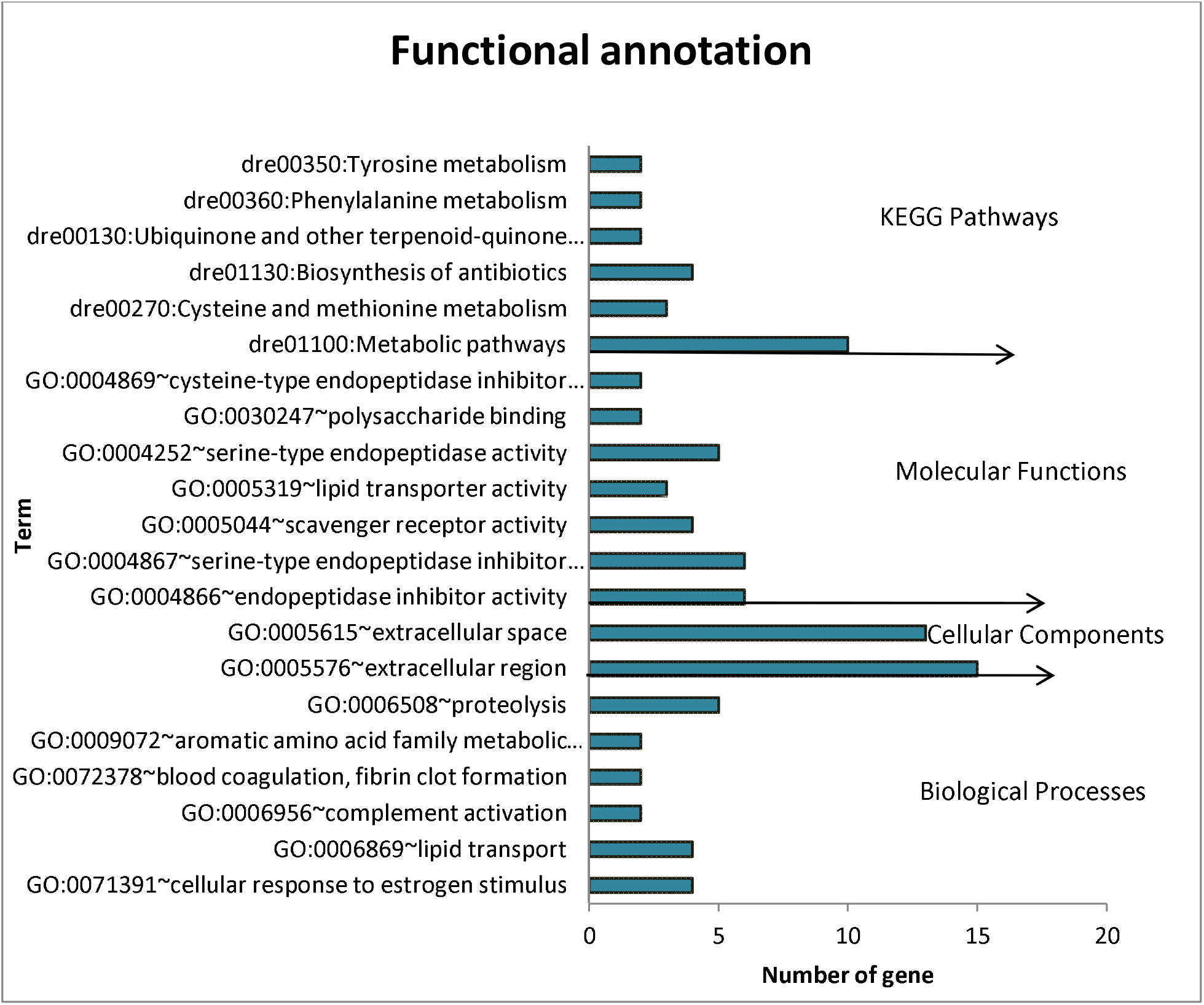
represents the most frequent GO terms and KEGG pathways, associated with our target genes. Biological processes, Cellular components), Molecular function, KEGG signaling pathways, where x-axis and y-axis show the number of genes and GO terms respectively and labeled with the p-value.

### 3.3. Network analysis and identification of hub genes

After removing the replicates the DEGs were submitted to the STRING database for PPI network construction, which involved 68 nodes and 202 edges with parameters including a minimum required interaction score > 0.4 (medium confidence). The whole PPI (Protein-Protein Interaction) network is graphically shown in **Figure 2**. MCODE analysis was divided the whole network in two clusters. In cluster i (**Figure 3**), the number of nodes involved were 14, numbers of edges involved were 90 and the MCODE score was 6.923. In cluster ii (**Figure 4**), the number of nodes involved were 10, numbers of edges involved were 48 and MCODE score was 5.333.

**Figure 2:**
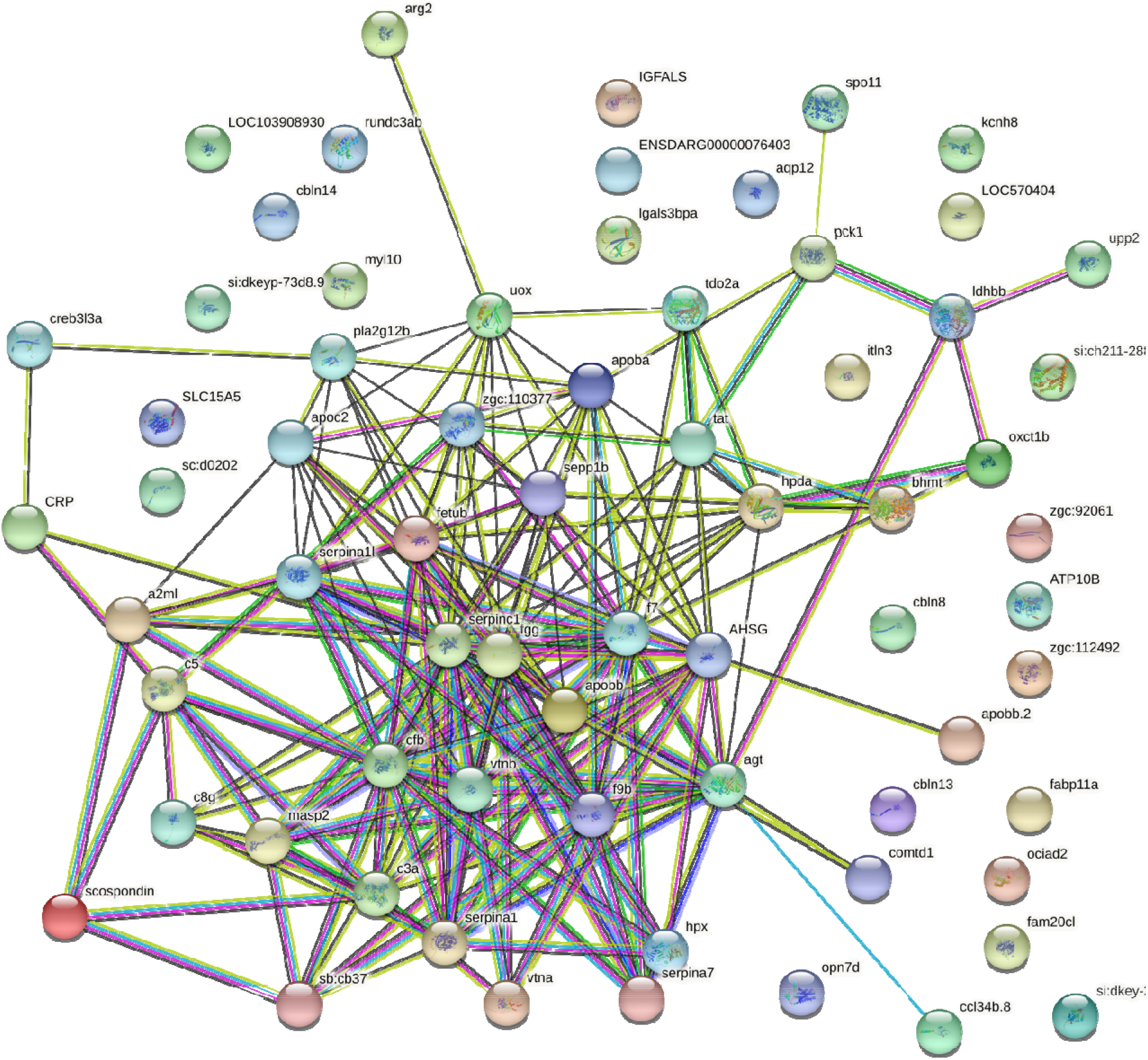
Interaction map of DEGs generated by STRING database.

**Figure 3:**
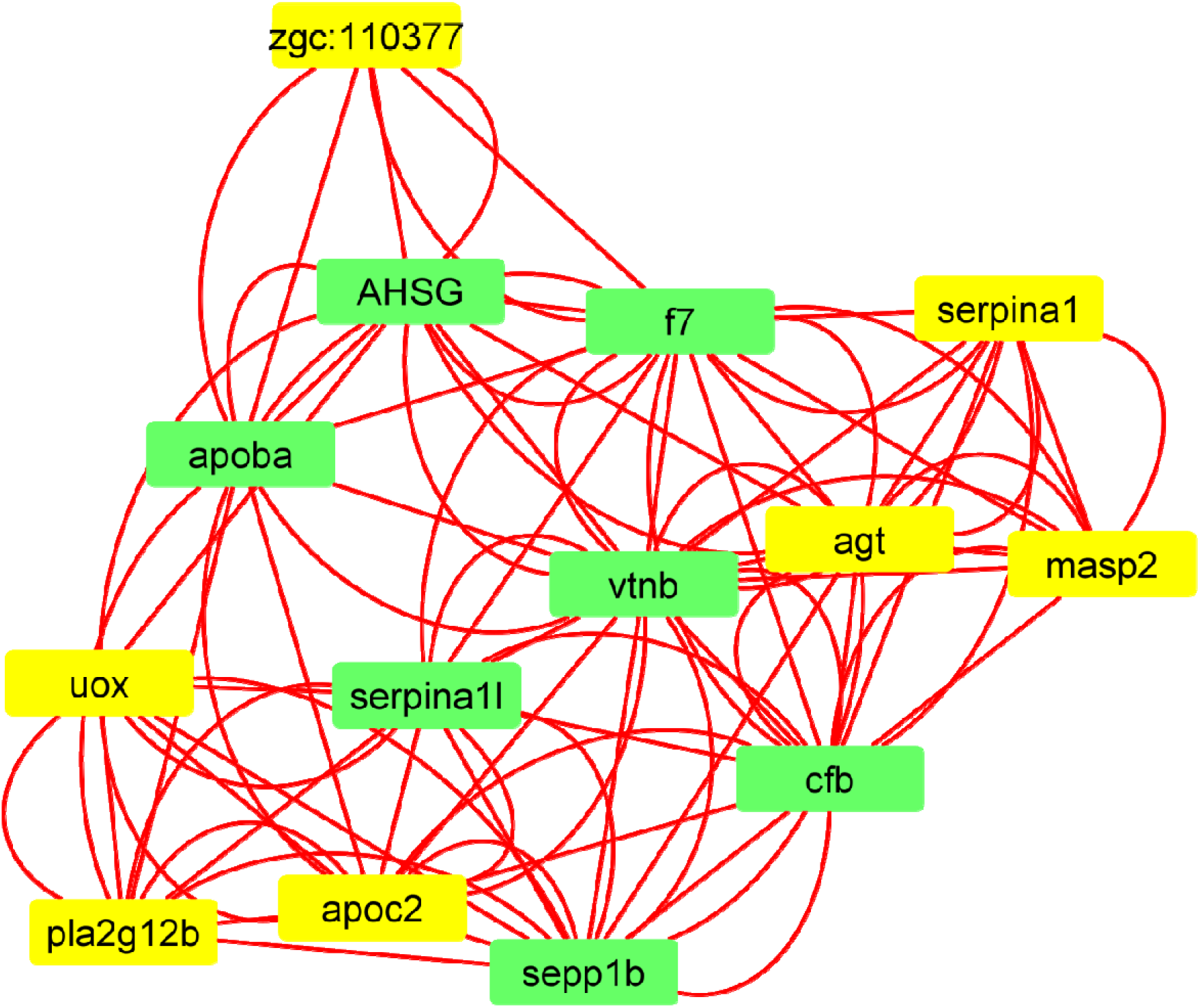
Various clusters predicted by Cytoscape on MCODE analysis. The colored nodes are those which are involved in the formation of that particular cluster in the network. The cluster i consists of 14 nodes and green colored nodes are hub nodes.

**Figure 4:**
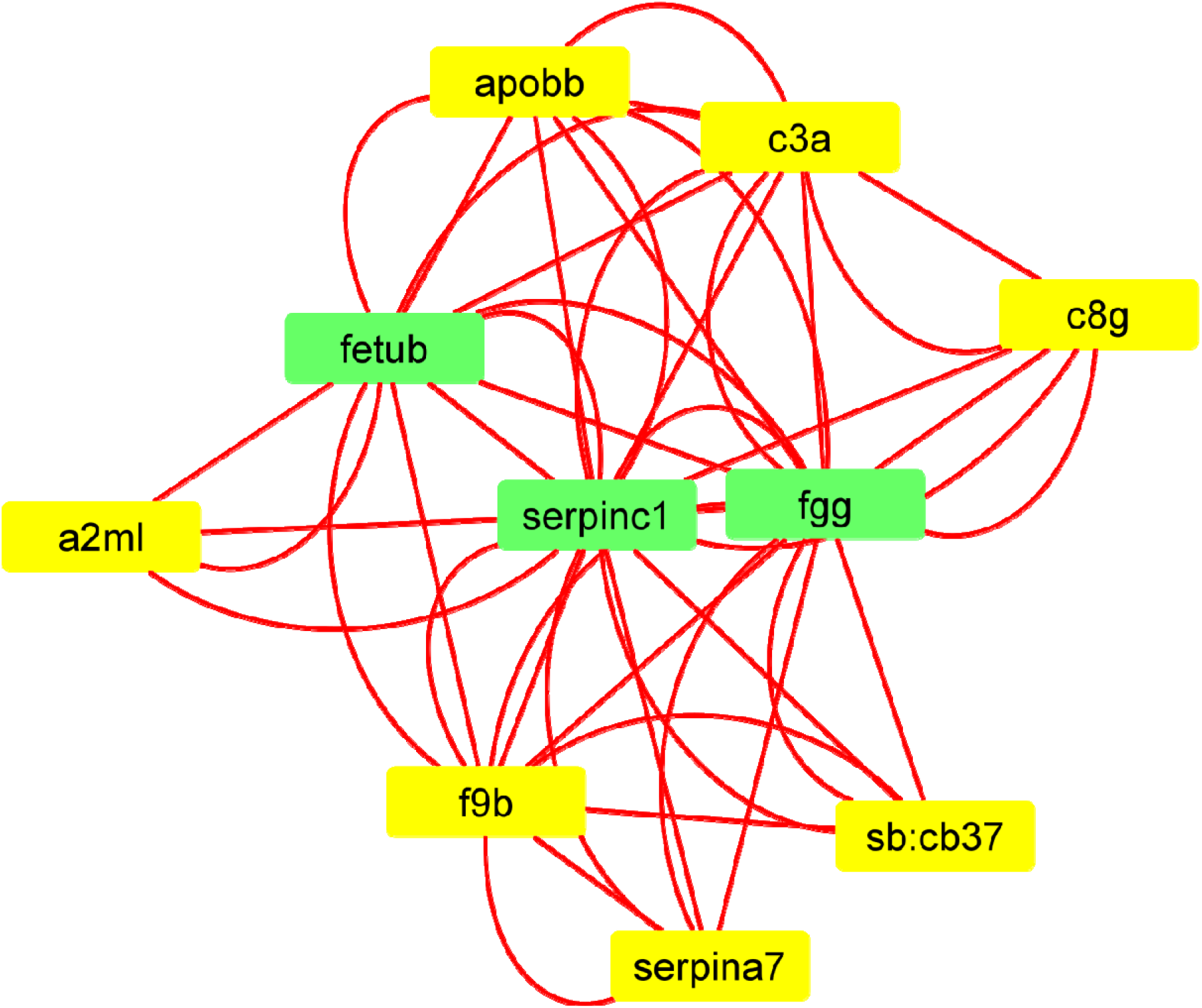
Various clusters predicted by Cytoscape on MCODE analysis. The colored nodes are those which are involved in the formation of that particular cluster in the network. The cluster ii consists of 10 nodes and green colored nodes are hub nodes.

From the Centiscape analysis, 1^st^ cluster (**Table 1**) predicted the top three bottleneck nodes like f7, vtnb and apoba, which having the high betweenness scores like 20.32727, 18.98788 and 9.733333 respectively. The top three hub nodes like vtnb, f7 and cfb are having high degree scores like 20, 18 and 16 respectively. The top three nodes with high closeness score were vtnb, f7 and cfb with scores like0.0625, 0.058824 and 0.055556 respectively. Hence, it was shown that the nodes may acts regulatory nodes by keeping other nodes together and influenced the activities of other nodes in this cluster as they having high betweenness, degree and closeness scores. In Cluster ii (**Table 2**), it was found that the top three nodes (serpinc1, fgg and fetub) were the bottleneck and hub nodes by having high scores of degree, betweenness and closeness at same time like (21.66667, 11.66667 and 5.333333), (18,16 and 12) and (0.111111, 0.1 and 0.083333) respectively, which may acts as the regulatory nodes for this cluster. Through the cytoHubba analysis in Cytoscape, the top ten hub genes were identified by the maximal clique centrality (MCC) method (**Figure 5**). The ten most significant genes were serpinc1, fgg, serpina1, apoc2, sepp1b, fetub, cfb, apobb and f7, where serpinc1 had the highest MCC score i.e. 32935 (**Table 3**). Hub nodes such as serpinc1, serpina1, apoc2 and fgg were found to be up regulated in infected zebrafish by comparing with uninfected zebrafish. Hub nodes such as sepp1b, fetub, cfb, apobb and f7 are found to be down regulated in infected zebrafish compared with uninfected zebrafish. The detailed information about these genes and their MCC scores with other network scores were listed in supplementary data Table S2 (**Page no 5-7**).

**Figure 5:**
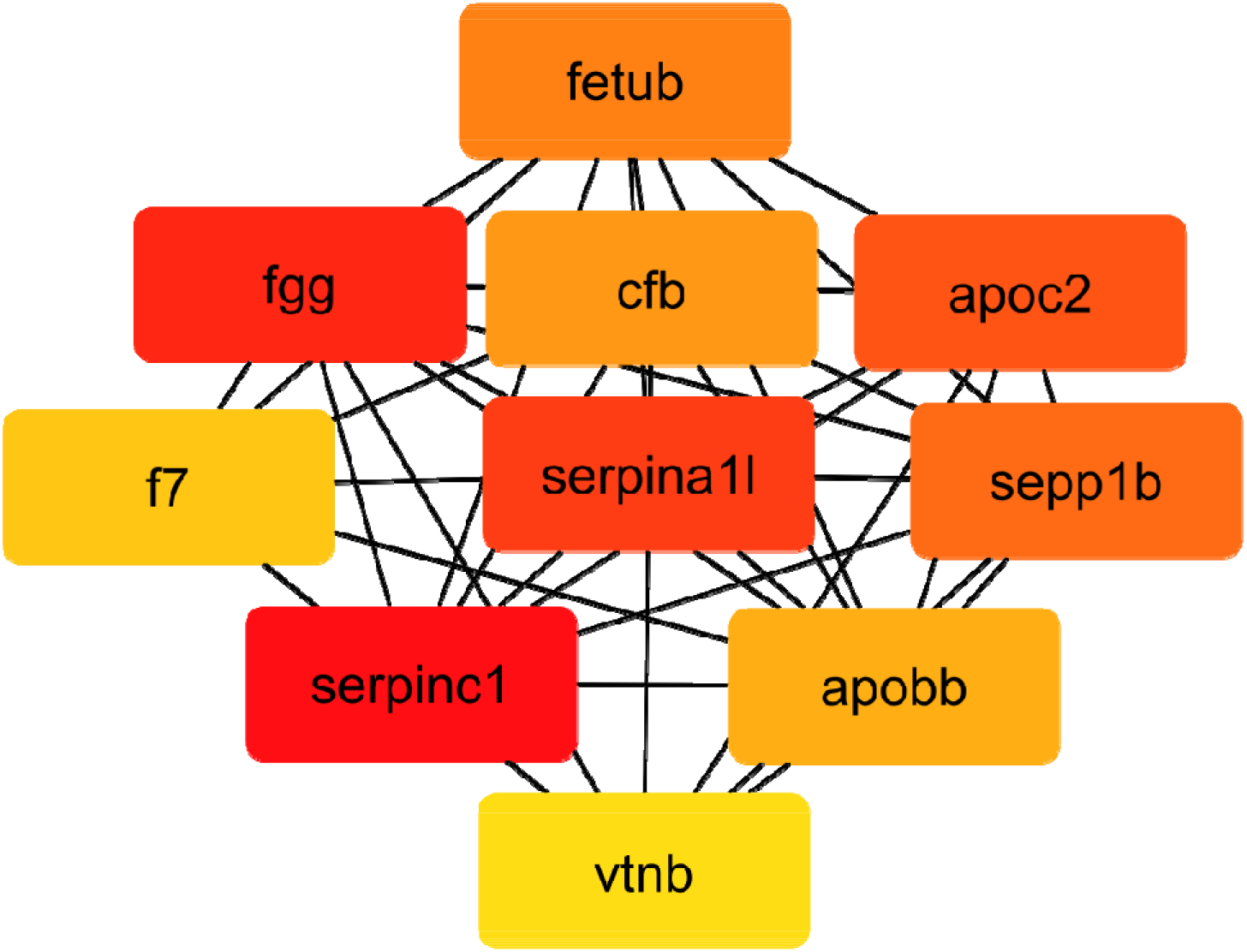
Top ten target genes based on highest MCC scores that were confirmed by CytoHubba analysis.

**Table 1:**
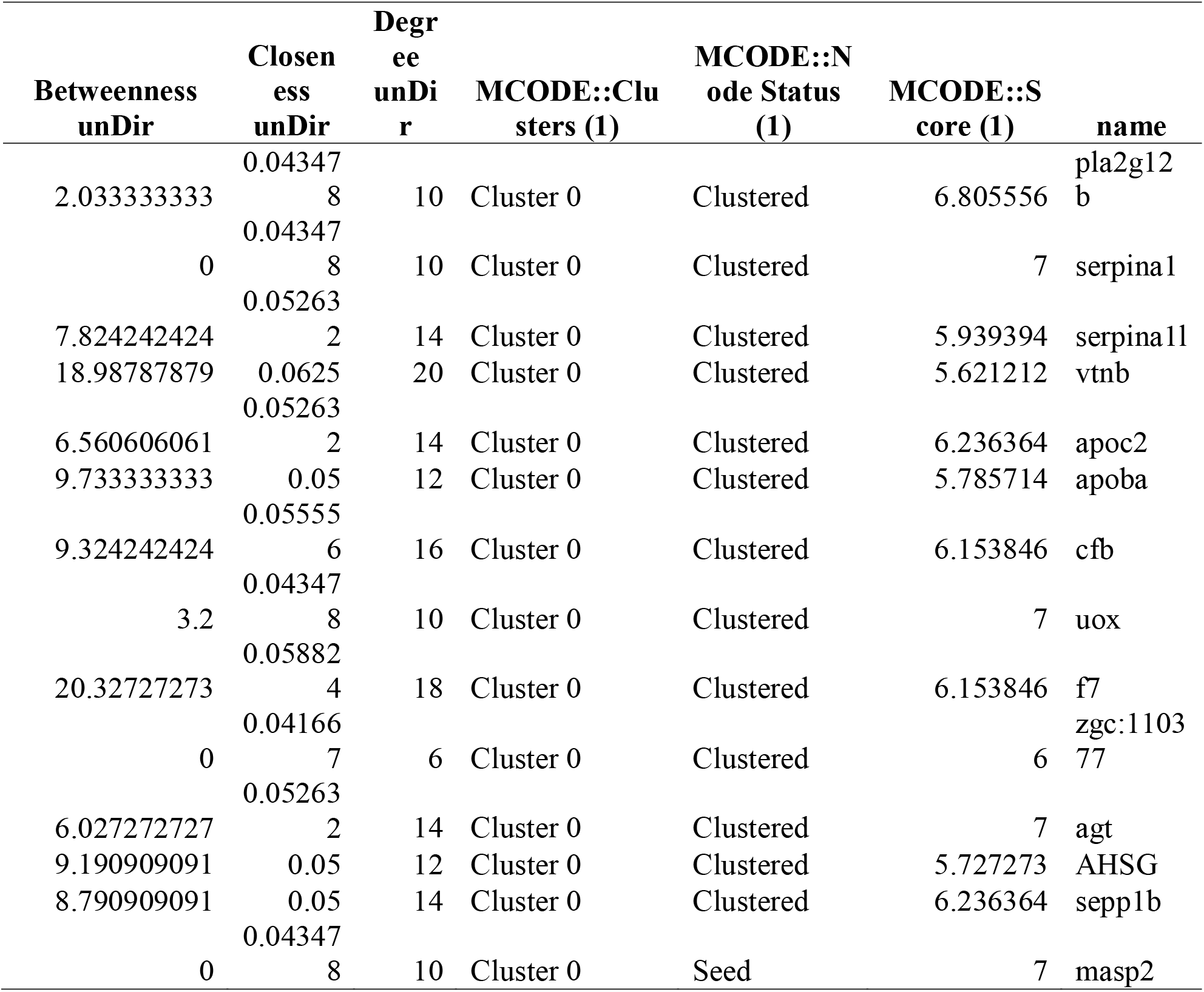
Centrality and cluster analysis of cluster i by MCODE and Centiscape

**Table 2:**
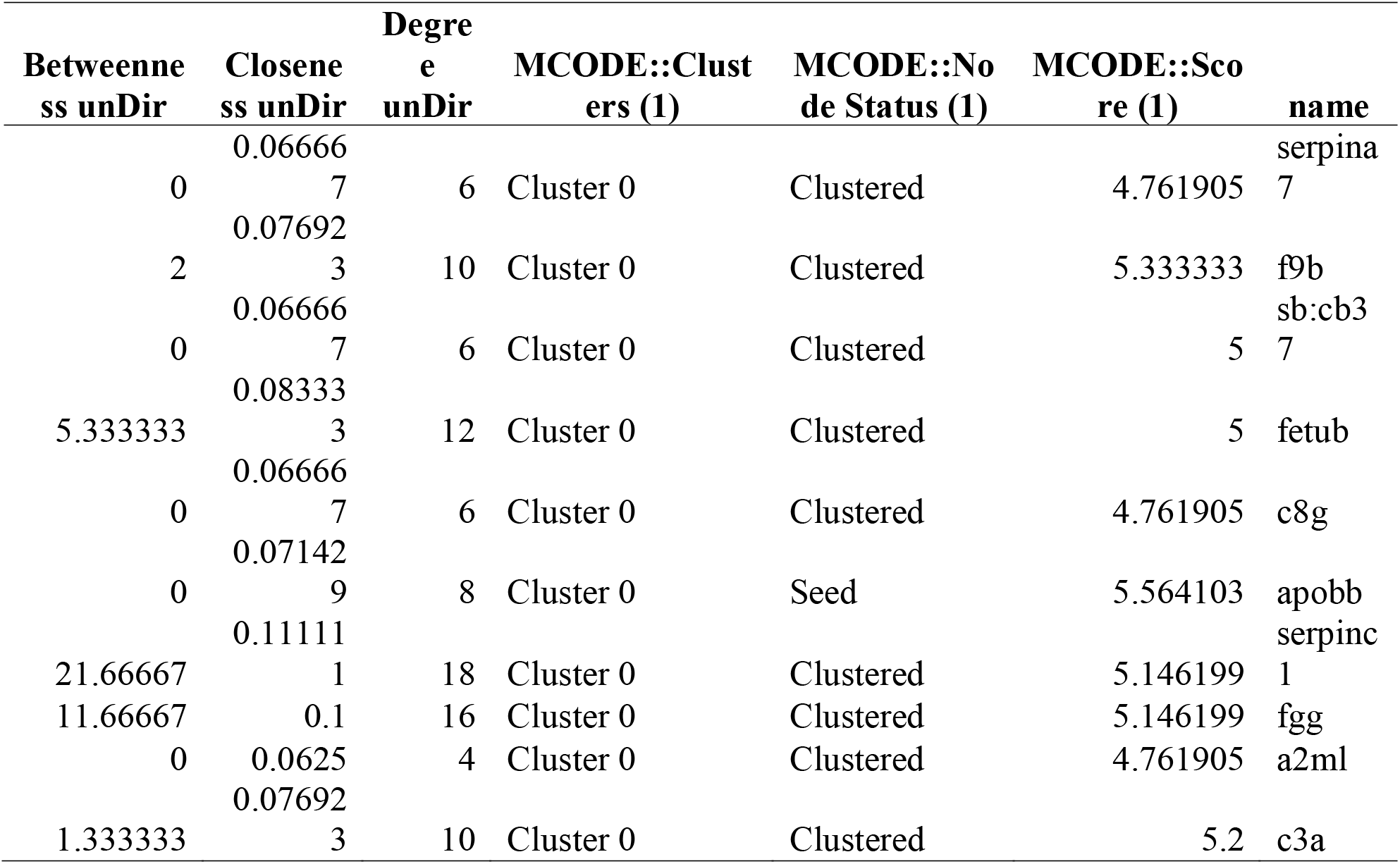
Centrality and cluster analysis of cluster ii by MCODE and Centiscape

**Table 3:** Top 10 hub nodes of predicted by CytoHubba

| Rank | Name | Score |
| --- | --- | --- |
| 1 | serpinc1 | 32935 |
| 2 | fgg | 32694 |
| 3 | serpina11 | 21960 |
| 4 | apoc2 | 20424 |
| 5 | sepp1b | 20281 |
| 6 | fetub | 19608 |
| 7 | cfb | 19254 |
| 8 | apobb | 18121 |
| 9 | f7 | 11852 |
| 10 | vtnb | 11334 |

Reported literatures are present on SVCV infection causing reduced coagulation cascade due to up regulation of inhibiting serpins like Serine proteinase inhibitor, clade C (serpinc1) and Serine proteinase inhibitor, clade a (serpinc1) serpina1 which favors SVCV-dependent host hemorrhages in Zebrafish [10, 23]. Lü *et al*. (2012) have reported enhanced expression complement factor B (cfb) in zebrafish upon infection with *Citrobacterfreundii*, where cfb genes significantly associated with skin immunity for complement activation [24]. The expression of the genes such as Serpina1, cfb, c8g and masp2 were found to be altered by transcription enhancement of the complement in the no presence of adaptive immunity. Some of the genes such as apoc2 and fgg have not been reported in SVCV infection and can be novel by our proposed network analysis approach as there is no reported literature.

## 4. Conclusion

This study analyzed the transcriptome between the infected and uninfected zebrafish with SVCV. 227 DEGs were screened respectively between the infected and uninfected zebrafish. The Gene ontology and pathway enrichment analysis were executed that aims to study the underlying mechanism of SVCV. Many DEGs were related to participate in the metabolic related pathways. In summary, the results obtained in our study establish the differential expression of hub genes and their related mechanism. These hub notes could act as potential biomarkers for SVCV infection. In conclusion, the expression pattern of hub genes, analyzed by our approach, may be considered as a molecular signature and could also offer potential biomarkers.

