## Supplementary Data for "Network analysis of Differentially Expressed Genes (DEGs) identified in zebrafish after infection with Spring viremia of carp virus (SVCV) – an *in silico* approach"

**Table S1: list of DEGs**

| ENSDARG00000091166 | ENSDARG00000051814 |
| --- | --- |
| ENSDARG00000040433 | ENSDARG00000062640 |
| ENSDARG00000045015 | ENSDARG00000078394 |
| ENSDARG00000023176 | ENSDARG00000002546 |
| ENSDARG00000035718 | ENSDARG00000067859 |
| ENSDARG00000074570 | ENSDARG00000013837 |
| ENSDARG00000018923 | ENSDARG00000040599 |
| ENSDARG00000040599 | ENSDARG00000037341 |
| ENSDARG00000068876 | ENSDARG00000068124 |
| ENSDARG00000002764 | ENSDARG00000075745 |
| ENSDARG00000071454 | ENSDARG00000090267 |
| ENSDARG00000051914 | ENSDARG00000062592 |
| ENSDARG00000077528 | ENSDARG00000076230 |
| ENSDARG00000077090 | ENSDARG00000078272 |
| ENSDARG00000074191 | ENSDARG00000089974 |
| ENSDARG00000038405 | ENSDARG00000093201 |
| ENSDARG00000020738 | ENSDARG00000092725 |
| ENSDARG00000022631 | ENSDARG00000043279 |
| ENSDARG00000056603 | ENSDARG00000013430 |
| ENSDARG00000078918 | ENSDARG00000053973 |
| ENSDARG00000069375 | ENSDARG00000039269 |
| ENSDARG00000086569 | ENSDARG00000019294 |
| ENSDARG00000010565 | ENSDARG00000093936 |
| ENSDARG00000069292 | ENSDARG00000090286 |
| ENSDARG00000089241 | ENSDARG00000034862 |
| ENSDARG00000093780 | ENSDARG00000056771 |
| ENSDARG00000076838 | ENSDARG00000018351 |
| ENSDARG00000054202 | ENSDARG00000042780 |
| ENSDARG00000004141 | ENSDARG00000056314 |
| ENSDARG00000053227 | ENSDARG00000090850 |
| ENSDARG00000092071 | ENSDARG00000096311 |
| ENSDARG00000035544 | ENSDARG00000044456 |
| ENSDARG00000096060 | ENSDARG00000037836 |
| ENSDARG00000004199 | ENSDARG00000015662 |
| ENSDARG00000030254 | ENSDARG00000038371 |
| ENSDARG00000043260 | ENSDARG00000003523 |
| ENSDARG00000090722 | ENSDARG00000042684 |
| ENSDARG00000057498 | ENSDARG00000060325 |
| ENSDARG00000079543 | ENSDARG00000070420 |
| ENSDARG00000020143 | ENSDARG00000093907 |
| ENSDARG00000009874 | ENSDARG00000094661 |
| ENSDARG00000012076 | ENSDARG00000016412 |
| ENSDARG00000046053 | ENSDARG00000055388 |
| ENSDARG00000089177 | ENSDARG00000076848 |
| ENSDARG00000088390 | ENSDARG00000037805 |
| ENSDARG00000045627 | ENSDARG00000088918 |
| ENSDARG00000088172 | ENSDARG00000075016 |
| ENSDARG00000030872 | ENSDARG00000037281 |
| ENSDARG00000041595 | ENSDARG00000070960 |
| ENSDARG00000021573 | ENSDARG00000045457 |
| ENSDARG00000036430 | ENSDARG00000093098 |
| ENSDARG00000013460 | ENSDARG00000022767 |
| ENSDARG00000079544 | ENSDARG00000032098 |
| ENSDARG00000038199 | ENSDARG00000053831 |
| ENSDARG00000011693 | ENSDARG00000012694 |
| ENSDARG00000036272 | ENSDARG00000078180 |
| ENSDARG00000069439 | ENSDARG00000052792 |
| ENSDARG00000020956 | ENSDARG00000087143 |
| ENSDARG00000063635 | ENSDARG00000056226 |
| ENSDARG00000076043 | ENSDARG00000054319 |
| ENSDARG00000038366 | ENSDARG00000088075 |
| ENSDARG00000070168 | ENSDARG00000079337 |
| ENSDARG00000019492 | ENSDARG00000041645 |
| ENSDARG00000070960 | ENSDARG00000021004 |
| ENSDARG00000007173 | ENSDARG00000092155 |
| ENSDARG00000022689 | ENSDARG00000075719 |
| ENSDARG00000042982 | ENSDARG00000069293 |
| ENSDARG00000019294 | ENSDARG00000007024 |
| ENSDARG00000093774 | ENSDARG00000069630 |
| ENSDARG00000042780 | ENSDARG00000007988 |
| ENSDARG00000015662 | ENSDARG00000090873 |
| ENSDARG00000039605 | ENSDARG00000055278 |
| ENSDARG00000032098 | ENSDARG00000038352 |
| ENSDARG00000079727 | ENSDARG00000078918 |
| ENSDARG00000094416 | ENSDARG00000029493 |
| ENSDARG00000024928 | ENSDARG00000060181 |
| ENSDARG00000018478 | ENSDARG00000038372 |
| ENSDARG00000053620 | ENSDARG00000024928 |
| ENSDARG00000059052 | ENSDARG00000036833 |
| ENSDARG00000029493 | ENSDARG00000037859 |
| ENSDARG00000077960 | ENSDARG00000078757 |
| ENSDARG00000088911 | ENSDARG00000090046 |
| ENSDARG00000088143 | ENSDARG00000071429 |
| ENSDARG00000052470 | ENSDARG00000026904 |
| ENSDARG00000059008 | ENSDARG00000077872 |
| ENSDARG00000070394 | ENSDARG00000013522 |
| ENSDARG00000076795 | ENSDARG00000053845 |
| ENSDARG00000063078 | ENSDARG00000071076 |
| ENSDARG00000079589 | ENSDARG00000055036 |
| ENSDARG00000069293 | ENSDARG00000052336 |
| ENSDARG00000076448 | ENSDARG00000017299 |
| ENSDARG00000060181 | ENSDARG00000012609 |
| ENSDARG00000089310 | ENSDARG00000076043 |
| ENSDARG00000042953 | ENSDARG00000079727 |
| ENSDARG00000004148 | ENSDARG00000086613 |
| ENSDARG00000089026 | ENSDARG00000045089 |
| ENSDARG00000070078 | ENSDARG00000000212 |
| ENSDARG00000038293 | ENSDARG00000094983 |
| ENSDARG00000024160 |  |
| ENSDARG00000079337 |  |
| ENSDARG00000076270 |  |
| ENSDARG00000090286 |  |
| ENSDARG00000007024 |  |
| ENSDARG00000042684 |  |
| ENSDARG00000013177 |  |
| ENSDARG00000075719 |  |
| ENSDARG00000090850 |  |
| ENSDARG00000038367 |  |
| ENSDARG00000018351 |  |
| ENSDARG00000037067 |  |
| ENSDARG00000091800 |  |
| ENSDARG00000094045 |  |
| ENSDARG00000026039 |  |
| ENSDARG00000075318 |  |
| ENSDARG00000037191 |  |
| ENSDARG00000094857 |  |
| ENSDARG00000041685 |  |
| ENSDARG00000056314 |  |
| ENSDARG00000086641 |  |
| ENSDARG00000070780 |  |
| ENSDARG00000094845 |  |
| ENSDARG00000019713 |  |
| ENSDARG00000059355 |  |
| ENSDARG00000045999 |  |
| ENSDARG00000093754 |  |
| ENSDARG00000091236 |  |
| ENSDARG00000059951 |  |
| ENSDARG00000078560 |  |
| ENSDARG00000052336 |  |
| ENSDARG00000035873 |  |
| ENSDARG00000073820 |  |
| ENSDARG00000090046 |  |

**Table S2: Showing the centrality measures**

| node_name | MCC | DMNC | MNC | Degree | EPC | BottleNeck | EcCentricity | Closeness | Radiality | Betweenness | Stress | ClusteringCoefficient |
| --- | --- | --- | --- | --- | --- | --- | --- | --- | --- | --- | --- | --- |
| spo11 | 1 | 0 | 1 | 2 | 1.188 | 1 | 0.2 | 13.23333 | 2.69048 | 0 | 0 | 0 |
| upp2 | 1 | 0 | 1 | 2 | 1.215 | 1 | 0.2 | 13.48333 | 2.7619 | 0 | 0 | 0 |
| oxct1b | 3 | 0.61557 | 2 | 6 | 2.774 | 1 | 0.25 | 19.75 | 3.80952 | 9.22967 | 528 | 0.13333 |
| c8g | 264 | 1.24398 | 7 | 14 | 6.183 | 1 | 0.25 | 22.33333 | 3.97619 | 1.95913 | 208 | 0.37363 |
| tat | 81 | 0.87472 | 8 | 18 | 6.046 | 4 | 0.33333 | 24.33333 | 4.19048 | 70.06249 | 5048 | 0.19608 |
| bhmt | 3 | 0.61557 | 2 | 6 | 3.046 | 1 | 0.25 | 18.58333 | 3.61905 | 1.47738 | 80 | 0.13333 |
| arg2 | 1 | 0 | 1 | 2 | 1.677 | 1 | 0.2 | 15.56667 | 3.19048 | 0 | 0 | 0 |
| pla2g12b | 10201 | 1.47972 | 9 | 20 | 7.159 | 2 | 0.25 | 24.58333 | 4.16667 | 67.27118 | 2688 | 0.32632 |
| pck1 | 4 | 0 | 1 | 8 | 2.652 | 2 | 0.25 | 19.16667 | 3.66667 | 95.37555 | 7216 | 0 |
| serpina7 | 240 | 1.33138 | 6 | 12 | 5.496 | 1 | 0.25 | 22.58333 | 4.07143 | 0.26873 | 24 | 0.42424 |
| comtd1 | 2 | 0 | 1 | 4 | 2.54 | 1 | 0.25 | 18.5 | 3.66667 | 0.84444 | 72 | 0 |
| ccl34b.8 | 1 | 0 | 1 | 2 | 1.786 | 1 | 0.25 | 17.08333 | 3.5 | 0 | 0 | 0 |
| sepp1b | 20281 | 1.45925 | 11 | 24 | 8.198 | 2 | 0.25 | 25.91667 | 4.2619 | 47.72174 | 2888 | 0.31159 |
| ldhbb | 4 | 0 | 1 | 8 | 2.491 | 2 | 0.25 | 19.66667 | 3.7381 | 102.6345 | 4720 | 0 |
| masp2 | 6150 | 1.32351 | 11 | 22 | 7.275 | 2 | 0.25 | 25.25 | 4.21429 | 11.75119 | 704 | 0.33766 |
| serpina1 | 5688 | 1.22934 | 12 | 24 | 7.813 | 2 | 0.25 | 25.75 | 4.2381 | 14.07332 | 656 | 0.30435 |
| hpda | 306 | 0.88234 | 11 | 22 | 7.474 | 1 | 0.33333 | 25.66667 | 4.28571 | 55.85793 | 3432 | 0.22511 |
| scospondin | 4 | 0.61557 | 2 | 8 | 3.293 | 1 | 0.2 | 19.2 | 3.61905 | 3.11318 | 56 | 0.07143 |
| serpina1l | 21960 | 1.37949 | 13 | 26 | 8.658 | 2 | 0.25 | 26.33333 | 4.2619 | 14.18403 | 800 | 0.33231 |
| apoc2 | 20424 | 1.40496 | 12 | 24 | 7.987 | 1 | 0.25 | 25.91667 | 4.2619 | 14.00447 | 1120 | 0.34783 |
| cfb | 19254 | 1.06856 | 20 | 40 | 9.98 | 3 | 0.25 | 29.91667 | 4.45238 | 76.85146 | 3240 | 0.22308 |
| a2ml | 241 | 1.33138 | 6 | 14 | 6.247 | 1 | 0.25 | 22.66667 | 4.02381 | 15.67947 | 1608 | 0.30769 |
| creb3l3a | 2 | 0 | 1 | 4 | 1.913 | 1 | 0.2 | 16.45 | 3.30952 | 1.07143 | 24 | 0 |
| c5 | 34 | 0.85588 | 6 | 16 | 5.185 | 2 | 0.25 | 22.41667 | 3.92857 | 22.93183 | 808 | 0.15 |
| CRP | 3 | 0 | 1 | 6 | 2.631 | 1 | 0.25 | 20.5 | 3.90476 | 17.80829 | 744 | 0 |
| zgc:110377 | 745 | 1.31715 | 7 | 16 | 6.477 | 1 | 0.25 | 23.75 | 4.14286 | 13.55947 | 976 | 0.3 |
| hpx | 30 | 1.03722 | 5 | 10 | 5.426 | 1 | 0.25 | 21.5 | 3.95238 | 0.84444 | 48 | 0.35556 |
| apobb | 18121 | 1.4635 | 12 | 26 | 8.707 | 2 | 0.25 | 26.33333 | 4.2619 | 42.11708 | 1824 | 0.30769 |
| apobb.2 | 1 | 0 | 1 | 2 | 1.838 | 1 | 0.25 | 17 | 3.47619 | 0 | 0 | 0 |
| tdo2a | 22 | 0.85588 | 6 | 12 | 4.961 | 1 | 0.33333 | 22.83333 | 4.11905 | 7.66032 | 640 | 0.27273 |
| f9b | 1416 | 1.07294 | 13 | 26 | 8.129 | 3 | 0.33333 | 26.66667 | 4.33333 | 26.98234 | 1616 | 0.25846 |
| vtna | 72 | 1.14118 | 6 | 12 | 5.622 | 1 | 0.25 | 22.16667 | 4 | 0.80096 | 56 | 0.36364 |
| fetub | 19608 | 1.19001 | 18 | 36 | 9.801 | 1 | 0.25 | 29.25 | 4.45238 | 47.96516 | 2768 | 0.25714 |
| vtnb | 11334 | 1.14169 | 15 | 30 | 9.077 | 1 | 0.33333 | 27.83333 | 4.40476 | 39.20818 | 2256 | 0.26207 |
| apoba | 1705 | 1.27697 | 10 | 22 | 7.539 | 1 | 0.33333 | 25.5 | 4.2619 | 90.79812 | 6568 | 0.27706 |
| serpinc1 | 32935 | 1.01763 | 27 | 56 | 11.709 | 5 | 0.33333 | 34.66667 | 4.7619 | 260.8109 | 9488 | 0.17922 |
| fgg | 32694 | 1.04575 | 26 | 52 | 11.265 | 1 | 0.33333 | 33.66667 | 4.71429 | 160.0662 | 7376 | 0.2006 |
| c3a | 2450 | 1.04739 | 13 | 26 | 8.525 | 1 | 0.25 | 26.41667 | 4.28571 | 54.85136 | 3528 | 0.25231 |
| sb:cb37 | 169 | 1.1708 | 7 | 16 | 6.114 | 1 | 0.25 | 23.16667 | 4.04762 | 17.86903 | 1712 | 0.26667 |
| uox | 5093 | 1.15725 | 10 | 22 | 7.357 | 2 | 0.25 | 25 | 4.16667 | 98.08777 | 5408 | 0.25108 |
| f7 | 11852 | 1.04003 | 21 | 42 | 10.326 | 2 | 0.33333 | 30.83333 | 4.54762 | 121.8968 | 4960 | 0.2137 |
| agt | 6735 | 1.17116 | 14 | 34 | 9.088 | 5 | 0.33333 | 29 | 4.47619 | 270.3827 | 9952 | 0.18538 |
| AHSG | 2895 | 1.00511 | 16 | 34 | 9.206 | 2 | 0.33333 | 28.83333 | 4.45238 | 137.9277 | 5168 | 0.19964 |
